# Comparative meta-proteomic analysis for the identification of novel plasmodesmata proteins and regulatory cues

**DOI:** 10.1101/2021.05.04.442592

**Authors:** Philip Kirk, Sam Amsbury, Liam German, Rocio Gaudioso-Pedraza, Yoselin Benitez-Alfonso

## Abstract

A major route for cell-to-cell signaling is via cell wall-embedded pores termed plasmodesmata (PD) forming the symplasm. PD regulate many aspects of plant development and responses to the environment however, our understanding of what factors affect their structure and permeability is limited. In this paper, a meta-analysis is presented as a tool for the identification of conditions affecting PD transport and *in silico* generation of PD proteomes for species of interest. The custom-built pipeline searches the whole genome for protein structural features and conserved domains identified on experimental proteomes and use it to predict PD candidates in 22 compatible plant species. Using the *in silico* proteome and microarray analysis, interactions between PD genes and conditions affecting PD function are identified. High salinity and osmotic stress affect a significant number of PD candidate genes and we provide evidence that these conditions regulate symplasmic transport of GFP. Using the pipeline, the *in silico* PD proteome for *Medicago truncatula* was generated, as an example of a plant in which experimental data is not available. The identification of a candidate receptor like protein was experimentally validated in *M. truncatula* transgenic roots expressing fluorescently tagged protein fusion. Together the results highlight the power of our newly designed tool in the identification of new factors and proteins influencing PD in diverse plant species.

## Introduction

Without careful coordination of the processes occurring in distant cells, plants would be incapable of responding effectively to changes in the environment, to the availability of resources for growth or to defend themselves against predators. Unsurprisingly, plants have evolved a myriad of long- and short- distance signalling pathways underpinning their ability to adapt and thrive in diverse conditions on Earth. A major route for signalling is the symplast, the continuous cytoplasmic connections established by cell walls-embedded pores termed plasmodesmata (PD). PD are dynamic structures tightly controlled to regulate intercellular signalling during development and in response to the environment (Li et al., 2021). Despite their importance, many questions remain unanswered especially regarding the molecular composition and the conditions that affect their function.

Over the last decade, proteomic analysis of PD-enriched membrane fractions has greatly improved our understanding of what proteins may localise to and associate with PD (Fernandez-Calvino et al., 2011; Park et al., 2017; Leijon et al., 2018; Brault et al., 2019). Localisation of labelled protein fusions using confocal, FRET-FLIM (Förster resonance energy transfer by fluorescence lifetime imaging) or transmission electron microscopy has experimentally confirmed PD association for around 60 proteins in the model plant *Arabidopsis thaliana* (Table S1). The PD proteomes include callose metabolic enzymes, such as callose synthase 3 (CALS3) and the PD-located β-(1,3)–glucanases PdBGs and AtBG_ppap (Sagi *et al.*, 2005; Vatén *et al.*, 2011; Benitez-Alfonso *et al.*, 2013). It also comprises signalling proteins and kinases such as the PD-Located Protein family (PDLPs) and the Lysin Motif domain-containing glycosylphosphatidylinositol-anchored protein 2 (LYM2) (Thomas *et al.*, 2008; Faulkner *et al.*, 2013). Proposed functions for many of these factors was recently reviewed in the context of cell-cell connectivity (Amsbury *et al.*, 2018; Dorokhov *et al.*, 2019).

Detection in PD proteomes is not proof of PD localisation as preparations of PD domains within cell walls are often contaminated with non-PD structures such as the plasma membrane (PM), cell walls and the endoplasmic reticulum (ER) (Brault *et al.*, 2019). The detection of proteins belonging to the ER is often attributed, not always verified, to the desmotubule (DT), a tubular structure that runs through PD connecting the ER of neighbouring cells. To eliminate contaminants, proteomic data from PM fractions is used but this is far from an ideal solution (Leijon et al., 2018; Brault et al., 2019). Contamination issues aside, the isolation of PD enriched fractions has been optimised for cell cultures which do not reflect the complexity and dynamic composition of PD associated with plant development and responses to the environment. Phylogenetic analysis can be used to identify family members expressed in distinct tissues but this is time consuming and predictions for diverse plant species is not always succesful.

In this paper, we present a meta-analysis to overcome some of the limitations imposed by proteomic screens. Comparing four published proteomes, we observed considerable overlap in protein subfamily composition. Based on this, we developed an analysis pipeline (implemented in R and named Plasmodesmata In silico Proteome 1 or PIP1; available as a resource) to rapidly generate candidate PD proteome lists for compatible species (currently 22 species). Overlap between predictions and experimental data demonstrates the validity of this approach. PD proteome lists were obtained *in silico* for the well-studied *A. thaliana* and for *Medicago truncatula*, a species for which a PD proteome has not been experimentally determined. Bioinformatic analysis of public transcriptomes allowed us to generate co-expression tables suggesting novel interactions between PD genes and conditions potentially affecting PD function. Using this approach and knowledge, we demonstrate the effect of salinity and high osmotic potential in the symplasmic unloading of GFP and confirm PD localization for a novel receptor like protein in *M. truncatula*. Together, the results highlight the power of our newly designed tool in the identification of new factors affecting PD in diverse plant species.

## Material and Methods

### Plant material and growth conditions

Seeds of *A. thaliana* plants expressing p*SUC2*::GFP were surface sterilised with ethanol. Control ATS media was prepared as described by Wilson et al. (1990) with 0.8% (w/v) agar (Type E, Sigma- Aldrich). When required, plates were prepared using ATS-based media supplemented with 3% (w/v) polyethylene glycol (MW 8000) or with 75 mM NaCl as indicated. Seeds were stratified at 4°C for 4 days before being transferred to long day light conditions (22°C, 16 h day, 150 mE/m2/s) for growth.

For *p35S::*Medtr1g073320-YFP transgenic roots, Gateway cloning was used to generate the vector following manufacture’s instruction (Invitrogen, USA). In brief, primers were designed to amplify Medtr1g073320 with linkers compatible for cloning into pDNR221 by BP reaction (Medtr1g0733201-Attb1: GGG GAC AAG TTT GTA CAA AAA AGC AGG CTC CAT GTT TTG ATT CTC TCT CCA; Medtr1g0733201-Attb2: GGG GAC CAC TTT GTA CAA GAA AGC TGG GTA CCA CAA ATC TCT TTC AGC CAA AA). Positive pDNR clones were confirmed by sequencing and used in LR reaction with the destination vector pB7YWG2 (Karimi *et al.*, 2002). The *p35S::*Medtr1g073320-YFP vector was amplified in *E. coli* and expressed in *Agrobacterium rhizogenes* for transformation.

Wild-type A17 *M. truncatula* seeds were lightly scarified with sandpaper and sterilised for 3 minutes in a 10% sodium hypochlorite solution, washed with water and left undisturbed in water for 4 hours. The seeds were transferred to agar-water plates and left in the dark for 7 days at 4 °C. Plates were transferred to RT overnight and *A. rhizogenes* carrying the *p35S-* Medtr1g073320-YFP was used for root transformation as described by (Boisson-Dernier *et al.*, 2001). After two to four weeks post-transformation, transgenic roots expressing YFP fusions were identified using fluorescent microscopy and selected for confocal imaging.

### PD proteome meta-analysis

PD proteomic data for *A. thaliana*, *Nicotiana benthamiana* and *Populus trichocarpa* were retrieved from their original publications (Fernandez-Calvino *et al.*, 2011; Park *et al.*, 2017; Leijon *et al.*, 2018; Brault *et al.*, 2019) and subfamilies annotated based on PANTHER16 (Mi *et al.*, 2021). The proteomes were incorporated into a pipeline using a custom-built R script (https://github.com/PhilPlantMan/PIP1). Instructions on how to use the pipeline are included in the GitHub repository along with instructions on how to customise pipeline parameters depending on user requirements. Necessary databases including the PD proteomes used here are packaged with the script.

The script dependencies include the R library ‘biomartr’ (Drost and Paszkowski, 2017). Protein features were predicted using the R library ‘ragp’ which integrates multiple tools for glycophosphatidylinositol anchor (GPI) and secretory signal peptide (SP) predictions (Emanuelsson *et al.*, 2007; Käll *et al.*, 2007; Pierleoni *et al.*, 2008; Almagro Armenteros *et al.*, 2019; Gíslason *et al.*, 2019; Dragićević *et al.*, 2020). Genes were classified as having a GPI and/or SP if at least one tool returned true for that feature. Transmembrane domains (TM) domains prediction was acquired from the ENSEMBL database annotation via the R package ‘biomartr’ using TMHMM (Krogh *et al.*, 2001). Predictions for N-myristoylation, S-farnesylation, S-geranylgeranylation, S-palmitoylation and S-nitrosylation were made using tools available by the Cuckoo workgroup (Xue *et al.*, 2010; Xie *et al.*, 2016; Ning *et al.*, 2020). For protein feature enrichment analysis, the tools described above were used to predict features for the whole *A. thaliana* genome (Araport11). Fisher’s exact test was used to determine statistical significance (cutoff provided in figure legends). To be fully pipeline-compatible for proteome input or to generate candidate gene output, the species must be listed in both Ensembl Plant databases (used by biomartr to retrieve sequence information) and in PANTHER16 (used to retrieve subfamily annotation) (Drost & Paszkowski, 2017; Howe *et al.*, 2020; Mi *et al.*, 2021). Currently, there are 22 compatible plant species (Table S2). For non compatible species, such as *N. benthamiana*, Arabidopsis orthologues can use. This enable integration of the PD proteome of Park et al. (2017) in the pipeline.

Lists of genes were compared by drawing a Venn or Euler diagram using the R library ‘eulerr’. The significance of the overlap between candidate lists and proteomes was determined using bootstrap analysis as follows. Sets of genes the same length as a candidate list and a proteome were randomly sampled from Araport11. The overlap in genes between the samples was recorded and repeated for *n* cycles (*n =* 10,000). Probability (*p*) was calculated as the proportion of cycles that attained an overlap at least as large as was observed between the candidate list and the proteome. The size of the overlap by chance was given as the median overlap in random samples over *n* cycles.

### Expression analysis

The gene correlation dataset ‘Ath-u.c1-0’ was downloaded from the ATTEDII database (Obayashi *et al.*, 2018). Optimal gene order and the corresponding dendrogram were computed using hierarchical clustering. The dendrogram was cut at an optimised height (h = 16) that gave a sufficient number/size of clusters (k = 151). These processes were performed in base R. Enrichment of genes within clusters from candidate lists and verified genes were determined using pairwise comparisons with Fisher’s exact test (p<0.05, holm-adjusted) via the R library ‘rcompanion ‘.

Publicly available microarrays for a subset of conditions were independently analysed. Microarray datasets (Affymetrix GeneChip Arabidopsis ATH1-121501) were downloaded from EBI ArrayExpress. For each experiment, expression data were normalised using the robust multi-array average (RMA) method and log_2_ transformed with the R package ‘oligo’ (Carvalho & Irizarry, 2010). Principal component analysis was used to identify and exclude outlier arrays and experiments with insufficient biological replicates. Genes with low levels of expression were filtered out. A design matrix was constructed for each experiment and a linear model applied using the R package ‘limma’ (Ritchie *et al.*, 2015). Differential expression of genes and a multiple comparison correction were determined using empirical bayes statistics via the package ‘limma’ and the results filtered by gene IDs. Heatmaps were constructed using the R package ‘ComplexHeatmap’ (Gu *et al.*, 2016).

### Confocal microscopy imaging for the analysis of GFP diffusion and protein localization

Seedlings of *A. thaliana* expressing *pSUC2:*:GFP were mounted on glass slides in 10 μg/ml propidium iodide (Sigma- Aldrich). Root tips were imaged using an LSM 800 upright confocal microscope (Zeiss, Germany). Profiles of fluorescence were determined using line and profiling tools in ImageJ. Lateral profiles were taken across the transition zone and rootward profiles started from the basal/apical meristem transition zone ending 150 μm towards the root tip. Fluorescence of lateral and rootward profiles were scaled between 0 and 1. Fluorescence across each lateral profile was binned (bins = 100) to compensate for small differences in root width. Fluorescence profiles of at least 6 plants per treatment were aggregated by calculating the mean (±SD) for position along the profile and plotted.

Transgenic roots expressing p*35S*::Medtr1g073320-YFP were stained with aniline blue fluorochrome (Biosupplies, Australia) in 0.1 M K_3_PO_4_ (pH 12) for callose detection. Roots were imaged by confocal microscopy (LSM-880, Zeiss) using 488 nm excitation laser for YFP and emission at 505–530 nm. Aniline blue was imaged with 405 nm excitation and emission at 463 nm.

## Results

### Workflow for the prediction of PD proteomes via meta-analysis of conserved subfamilies and structural domains

Proteomic data obtained from *A. thaliana* (Fernandez-Calvino *et al.*, 2011; Brault *et al.*, 2019), *N. benthamiana* (Park *et al.*, 2017) and *P. trichocarpa* (Leijon *et al.*, 2018) PD-enriched fractions were fed into a custom-built R-based pipeline released as a resource with this article (https://github.com/PhilPlantMan/PIP1). The workflow is described in Figure 1 and in the Material and Methods section. First, subfamilies of genes identified in at least one of the PD proteomes mentioned above were annotated using PANTHER16. Subfamilies classifications are used to extract all genes making the *in silico* proteome for a particular plant of interest. For some applications, using families classifications may be more appropriate and PIP1 prompts users whether to generate candidates based on family or subfamily identifiers. Data presented in this paper are based on candidate generation using subfamily identifiers as a more restrictive approach leading to shorter candidate lists (see example for Arabidopsis below). The pipeline categorises the output by whether the subfamily is present in one or multiple PD proteomes and based on predictions of distinctive features identified in PD verified genes (Figure 1). Features of membrane-bound proteins were predicted for the list of PD verified genes in Table S1 using multiple online platforms as described in Material and Methods. When compared to the Arabidopsis whole proteome, PD-localised proteins are overrepresented in predicted GPI anchors, s-geranylgeranylation, s-palmitoylation, SP and TM domains (Figure S1). The presence of a SP in combination with either a GPI and/or TM domain returned the highest proportion of verified genes thus these features were chosen for gene categorisation.

**Figure 1.**
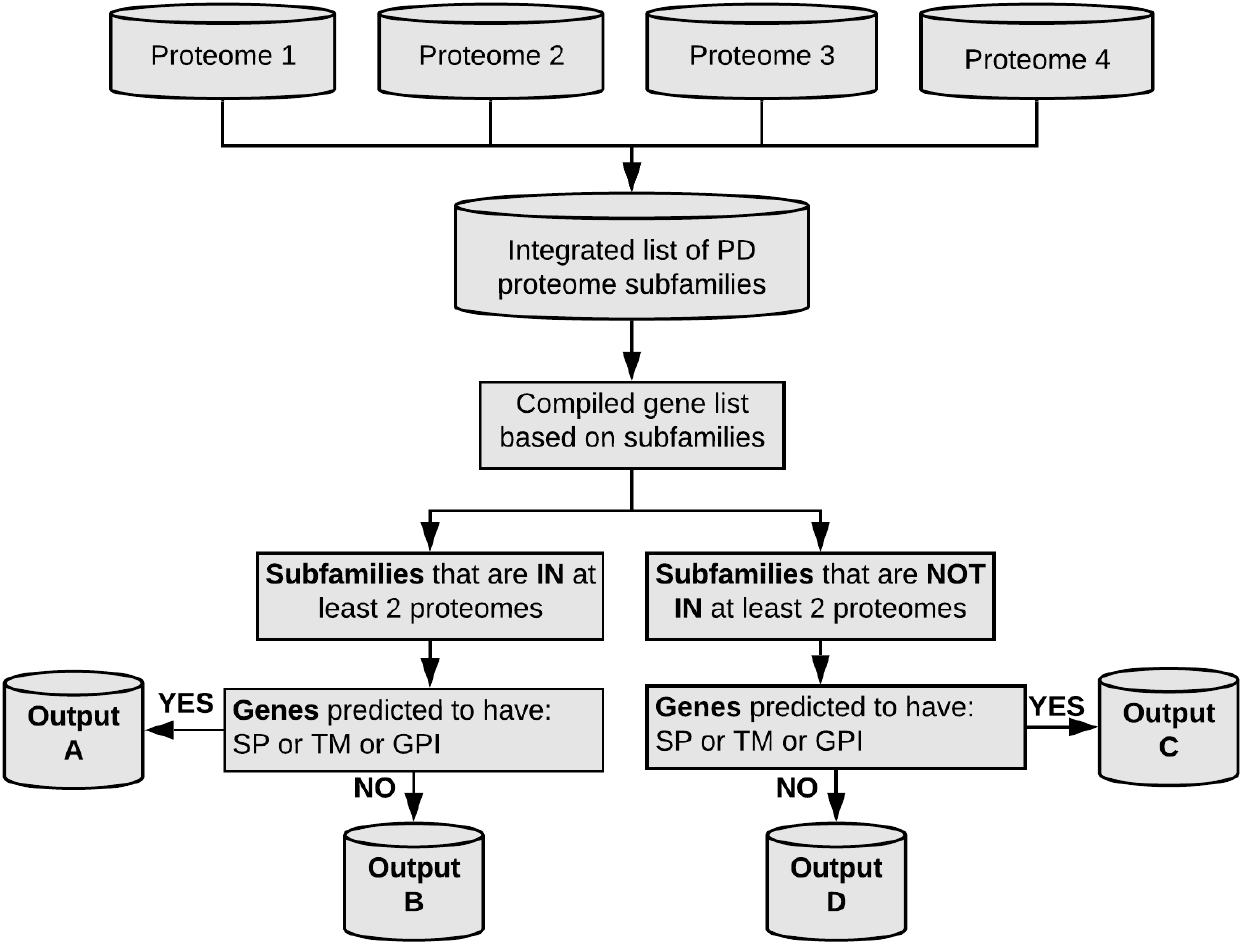
Pipeline to determine candidate plasmodesmata (PD) proteome. 4 proteome databases have been determined experimentally by Fernandez-Calvino et al., 2011; Park et al., 2017; Brault et al., 2019 and Leijon et al., 2018. A list was compiled of subfamilies of proteins identified in the different experimental proteome databases using PANTHER16. Genes belonging to these subfamilies were extracted for the target plant species. These were classified by whether its subfamily was present in at least two PD experimental proteomes or just one. Further classification was based on whether genes were predicted to have a signal peptide (SP), transmembrane domain (TM) or glycosylphosphatidylinositol anchor (GPI) as these features are enriched in verified PD genes (Figure S1). As a result, 4 list of genes were obtained (A-D) representing the predicted PD proteome for the target plant.

As a result, the pipeline outputs 4 lists of genes: list A are genes of subfamilies present in more than one PD proteome containing either a SP, GPI or TM domains; list B are also present in multiple proteomes but lacking the membrane localising features, list C and D are genes in subfamilies found in a single proteome either with (list C) and without (list D) the predicted SP, GPI or TM features (Figure 1). Using this approach, 1882 gene subfamilies were identified in at least one PD proteome (Table S3). As expected, there is substantial overlap between subfamilies identified in the different proteomes, with 311 subfamilies identified in more than one proteome (Table S3), forming list A and B. As PD verified genes usually display membrane targeting features, list A should be considered the priority list but, since only four proteomes are available, list C cannot be discarded. In support of this prediction, list A contains subfamilies with known PD activities such as enzymes involved in callose metabolism (BG and CALS) and receptors involved in signalling (such as PDLPs).

### Applying PIP1 to proteome predictions in *Arabidopsis thaliana* and poplar

For *Arabidopsis*, the *in silico* proteome comprises list A with 206 genes (158 subfamilies), list B: 208 genes (152 subfamilies), list C: 751 genes (597 subfamilies) and D: 1117 genes (802 subfamilies) (Table S4). Most known PD genes are isolated from studies in *A. thaliana*. To validate our approach, we determined the number of PD verified genes identified by the pipeline. Generated candidate lists contain 60.7% of the genes known to localise at PD identified either using fluorescent translational fusions and/or isolated in independent, non-proteomic studies (compare Tables S4 and S1). These include the beta-1,3 glucanase AtBG_ppap and C2 calcium/lipid-binding plant phosphoribosyltransferase family protein MTCP6 and MTCP9 identified in the Brault *et al.* (2019) proteome and proteins identified in independent studies such as the leucine-rich repeat receptor-like serine/threonine-protein kinase BAM2 (Rosas-Diaz *et al.*, 2018). We failed to output known PD proteins such as CRINKLY4 (ACR4) or the PD-located protein 5 (PDLP5) likely because these subfamilies are low expressed or absent in cell cultures. Extending the output to family members instead of just subfamily members increased the coverage of known PD genes to 83.6% but consequently increased the total candidate gene size to over 9000 genes (Table 1). Genes in list A and C included in families containing at least one previously verified PD-localised protein were identified as the most likely targets for future experiments (Table S5). In Arabidopsis, this list comprises 55 new proteins.

**Table 1.**
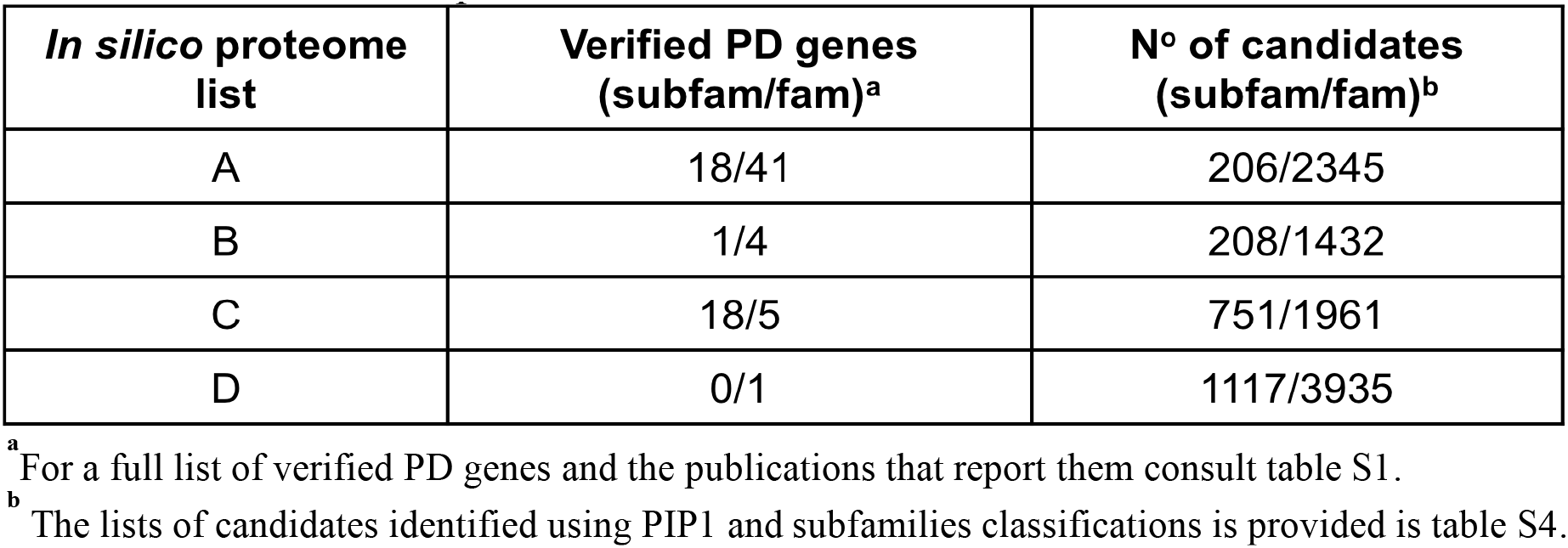
Verified PD genes and candidates identified in the Arabidopsis *in silico* proteome. The number of genes identified when using either subfamilies (subfam) or families (fam) classifications in PIP1 is provided.

| ***In silico* proteome list** | **Verified PD genes (subfam/fam)<sup>a</sup>** | **N° of candidates (subfam/fam)<sup>b</sup>** |
| --- | --- | --- |
| A | 18/41 | 206/2345 |
| B | 1/4 | 208/1432 |
| C | 18/5 | 751/1961 |
| D | 0/1 | 1117/3935 |

<sup>a</sup> For a full list of verified PD genes and the publications that report them consult table S1.
<sup>b</sup> The lists of candidates identified using PIP1 and subfamilies classifications is provided in table S4.

Despite lacking features for PD localisation, list B contains a considerable number of genes identified in more than one PD proteome. The number of known PD proteins identified in list B and D suggest they might contain contaminants but it could also contain PD proteins with no predicted membrane targeting domains such as Remorin 2 REM1.2 (Huang *et al.*, 2019). We explored the hypothesis that list B might contain mobile proteins unintentionally captured within PD fractions. To test this hypothesis, the candidate PD gene lists were compared to the mobile proteome identified in *Cuscuta australis* (dodder) parasitising *A. thaliana* (Liu *et al.*, 2020). Candidate mobile proteins were identified in all candidate lists but list B had the largest overrepresentation (20.2%) with an overlap over 10x greater than would be expected by chance (Figure S2).

To evaluate the strength of the pipeline further, we generated candidate PD lists for *P. trichorcarpa*. 1032 out of 1148 genes/proteins identified in the PD-enriched fraction of poplar cell cultures (Leijon *et al.*, 2018) were predicted in the candidate lists generated with our pipeline. The lists contained 182 (list A), 129 (list B), 397 (list C) and 289 (list D) of the genes identified in the PD-enriched fraction (Leijon *et al.*, 2018). For list A and C the *in silico* proteome is 50% larger than the experimental proteome, reflecting inclusion of proteins not expressed in cell cultures. 116 genes identified in the experimental proteome are excluded because they are not annotated within PANTHER subfamilies. When the poplar proteome is excluded as input in the pipeline, the overlap between the candidates lists and the poplar PD proteome remains high; 20x larger than would be expected by chance significant (Figure S3). In summary, our results validate the use of PIP1 as a new resource to generate candidate PD genes, and potentially mobile proteins, without the cost of time and resources associated with growing and maintaining cell cultures carrying out their biochemical isolations, fractionations and protein sequencing. The *in silico* proteome list is larger than experimental proteomes because there are not constrictions associated with protein expression in cell cultures.

### Drought and salinity regulate the expression of PD candidates and symplasmic transport

The generation of PD proteomes not only enables target gene selection but, in combination with transcriptomics, can reveal conditions that regulate PD genes. To establish whether this is the case, co-expression data extracted from ATTED-II database was ordered by hierarchical clustering to identify co-expressed genes in the *in silico* proteomes. For *A. thaliana*, two clusters were found to be significantly over-represented (relative to the wider ATTED-II gene set) in PD-verified genes: cluster 87 and cluster 100 (Figure 2 A, B). 11 genes from list A and 18 from list C were found in cluster 87, whereas cluster 100 contained 16 genes of list C (see Table S5 for examples). The expression of PD-candidates in clusters 87 and 100 was also analysed using publicly available microarrays as described in Materials and Methods. Figure 2C and Figure S4 show that the expression of the candidate genes overlaps with PD-marker proteins such as AtBG_ppap, PDCB (PD callose binding proteins) and PDLP family members.

**Figure 2:**
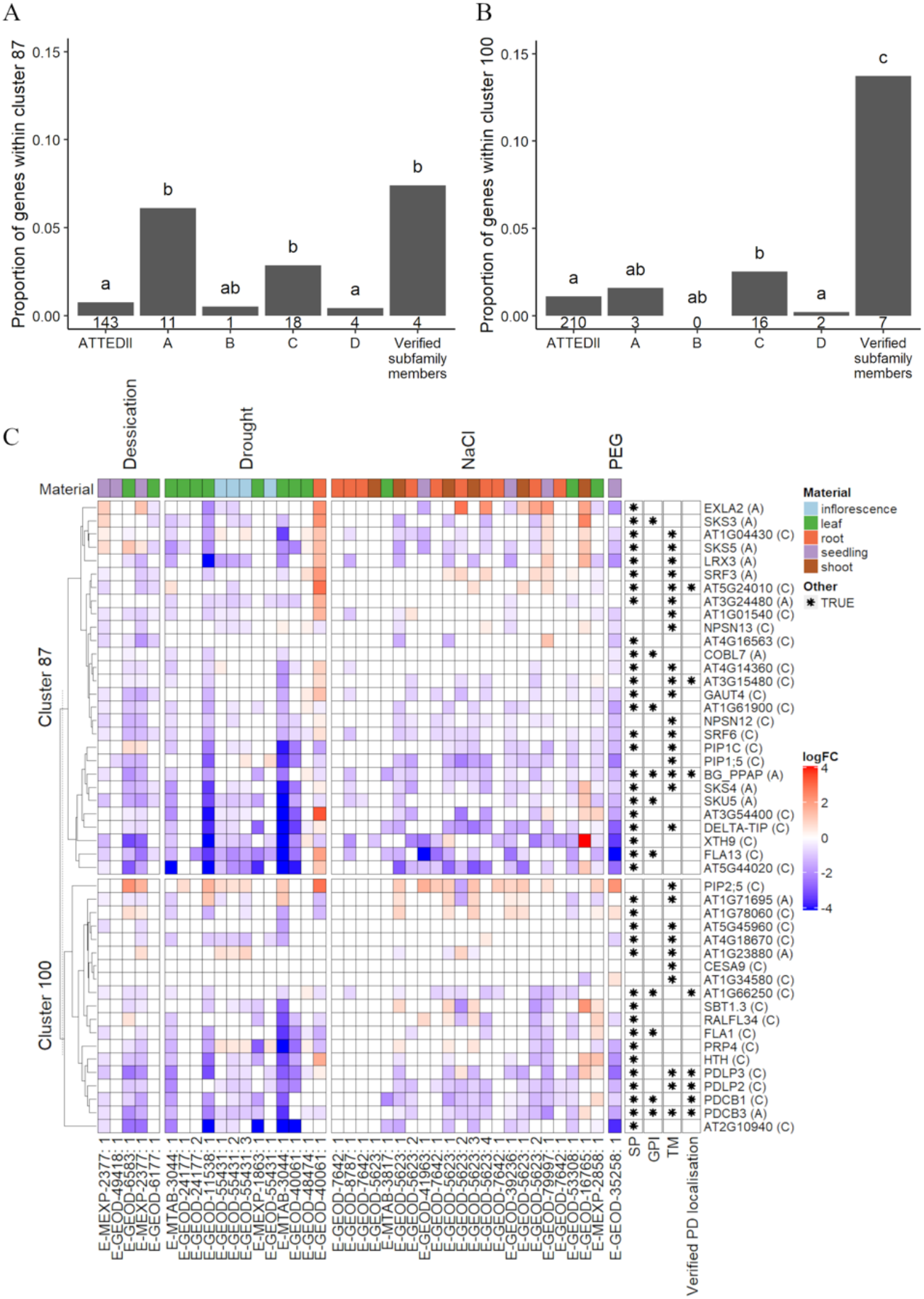
Expression analysis of PD candidates and PD verified genes. *Arabidopsis thaliana* co-expression data from ATTEDII were subset into clusters based on hierarchical clustering (k = 151). Representation analysis of the total ATTEDII database, candidate lists A-D and verified PD subfamilies was performed on each cluster; (A) cluster 87 and (B) cluster 100 show significant overepresentation in verified subfamily members. Number underneath each bar = number of genes within that cluster. Bars with differing letters above are significantly different (Fisher’s exact test, Holm corrected, p ≤ 0.05). (C) Differential expression (log_2_FC, ranges from blue: downregulated to red: upregulated in relation to control) of genes in clusters 87 and 100 (only list A, C and verified genes are shown) determined using public microarrays from experiments pertaining to osmotic stress. Other treatments are presented in Supplemental figure 4. Columns labels bottom = ArrayExpress accession codes followed by a reference number. See Supplemental table 6 for further information. (left) Rows are grouped by cluster and then (dendrogram) ordered by hierarchical clustering. (right) Gene ID and properties. Asterisks in cells denote predictions on membrane targeting features (e.g. is predicted to have a SP). SP = Signal peptide, GPI = glycosylphosphatidylinositol anchor, TM = transmembrane domain. Cell colour above each column = plant material sampled.

When comparing the expression of PD-verified and PD-candidate genes in a variety of biotic and abiotic stress conditions (Figure S4), osmotic stress (drought, desiccation, elevated NaCl and exogenous treatment with PEG) was found to be a strong regulator (Figure 2C). To determine if these changes in expression reflect conditions affecting symplasmic transport, cell-to-cell connectivity was tested in root tips of transgenic *A. thaliana* seedlings expressing cytosolic GFP, driven by the phloem companion cell specific promoter *Sucrose Symporter 2 SUC2* (p*SUC2*::GFP) (Imlau *et al.*, 1999).

p*SUC2*::GFP seeds were germinated and grown in control conditions, 3% PEG and 75mM NaCl (Hunter *et al.*, 2019). Media water potential (at 25 °C) was estimated by adding the estimated solute potentials of individual medium components based on empirical and modelled data (Robinson & Stokes, 1949; Ghashghaie *et al.*, 1991; Gopal & Iwama, 2007). The water potential was reduced in both 3% PEG and 75mM NaCl (Figure 3A). Roots were imaged using confocal microscopy and both lateral and rootward diffusion of GFP from phloem companion cells were measured using mean grey values in Image J. The short and long term effects of changes in water potential were evaluated by comparing changes in GFP distribution in roots grown for 4 days directly in PEG and NaCl media (Figure 3A) and roots grown for 4 days in control media and transferred for 24h to PEG and NaCl media (Figure 3D). There were no noticeable changes in lateral distribution of GFP across primary roots grown permanently in PEG and NaCl relative to control conditions (Figure 3 A, B) but the rootward profiles, show a decrease in fluorescence in both, 3% PEG and 75mM NaCl (Figure 3C). The effect is stronger in NaCl rootward fluorescence profiles, particularly between 50 μm - 150 μm towards the tip where relative fluorescence drops over 13% in 75 mM NaCl and less than 2% in control conditions (Figure 3C).

**Figure 3:**
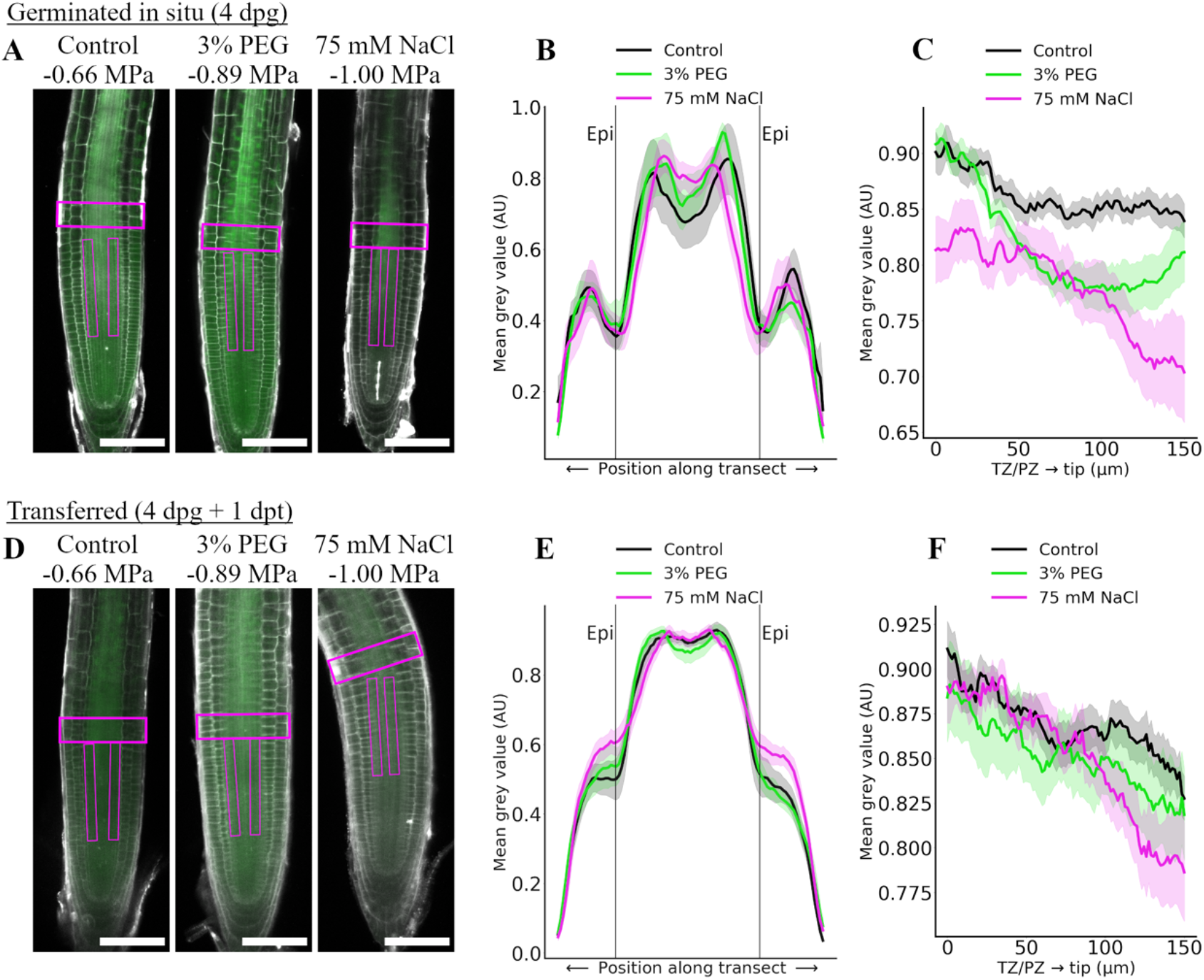
GFP diffusion in the root meristem of *Arabidopsis thaliana* exposed to PEG and sodium chloride. (A) Seeds expressing p*SUC2*::GFP were grown on ATS control media, ATS supplemented with 3% (w/v) polyethylene glycol (PEG) or with 75 mM NaCl. Roots were counterstained with propidium iodide (PI) and images sequentially collected 4 days post-germination (4 dpg) at 561 nm (PI shown in grey) and 488 nm (green: GFP). Estimated water potential (MPa, see main text) is labelled on top of the representative pictures. (B-C) Transects highlighted by magenta boxes in A were used to determine GFP fluorescence profiles using mean grey values (AU: arbitrary units). (B) shows transects across the transition zone (TZ) while (C) shows transects rootward starting from the transition/proliferation zone (TZ/PZ) towards the root tip. (D-F) Similar experiments were carried out with seedlings germinated on ATS media and transferred for 1 day (1dpt) to 3% PEG or 75 mM NaCl. Scale bar = 100 μm. Charts show traces representing min-max normalised GFP fluorescence along the transect for N>6 plants (mean ± SE).

No significant differences in GFP distribution were observed 24 hours after transfer from control to 3% PEG conditions (Figure 3 D-F) but fluorescence was diminished in the rootward profile of roots transferred to 75 mM NaCl (Figure 3F). Over the length of 150 μm from the tip, relative fluorescence decreased 11% in plants transferred to 75 mM NaCl. Lateral profiles in roots transferred to 75 mM NaCl show a higher fluorescence (likely auto-fluorescence) in the epidermis (Figure 3E).

Taken together, these data suggest that regulation of PD candidates in osmotic stress is translated into changes in symplasmic connectivity demonstrating the potential for using combined proteomic and transcriptomic meta-analysis as a method to determine conditions regulating PD function and therefore symplasmic communication.

### A meta-analysis approach predicts plasmodesmata proteins in *Medicago truncatula*

One of the advantages to the pipeline outlined here is that it can be applied to species for which an experimentally determined PD proteome is not publicly available. To demonstrate this, the pipeline was used to generate candidate PD gene lists for *M. truncatula* (Table S6). List A and C comprised 1018 genes belonging to subfamilies represented in at least one experimental proteome and displaying membrane targeting features (SP, GPI or TM). These lists include *M. truncatula* orthologues of the CALS, PDCB and PDLP genes (Table S7).

Transcriptomic profiling was used to determine clusters of genes co-regulated within the predicted proteome lists A and C. Published microarray data from early rhizobia inoculation, various nitrate concentrations and nodulating roots (Table S8) were re-analysed. These conditions were selected as they have been previously shown to regulate PD genes and symplasmic communication (Haney & Long, 2010; Crook *et al.*, 2016; Gaudioso-Pedraza *et al.*, 2018). The β-(1,3)-glucanase, MtBG2 (Gaudioso-Pedraza et al., 2018), and the Super Numeric Nodule receptor (SUNN, Crook *et al*., 2016) are, to our knowledge, the only verified PD-localised proteins in *M. truncatula*. A cluster containing MtBG2 and SUNN was identified comprising 70 genes from list A and C (Figure 4). Genes in this cluster share a similar expression pattern, particularly in roots post-inoculated with rhizobia (E-MEXP-1097). Interestingly, four genes encoding receptor-like proteins show a strikingly similar expression profile to MtBG2. These are MTR_5g083910, MTR_4g014070, MTR_1g073320 and MTR_2g011180. MTR_1g073320 (also known as Medtr1g073320) was selected for further studies because it contains sequence domains characteristic of PDLP family members (Table S7).

**Figure 4:**
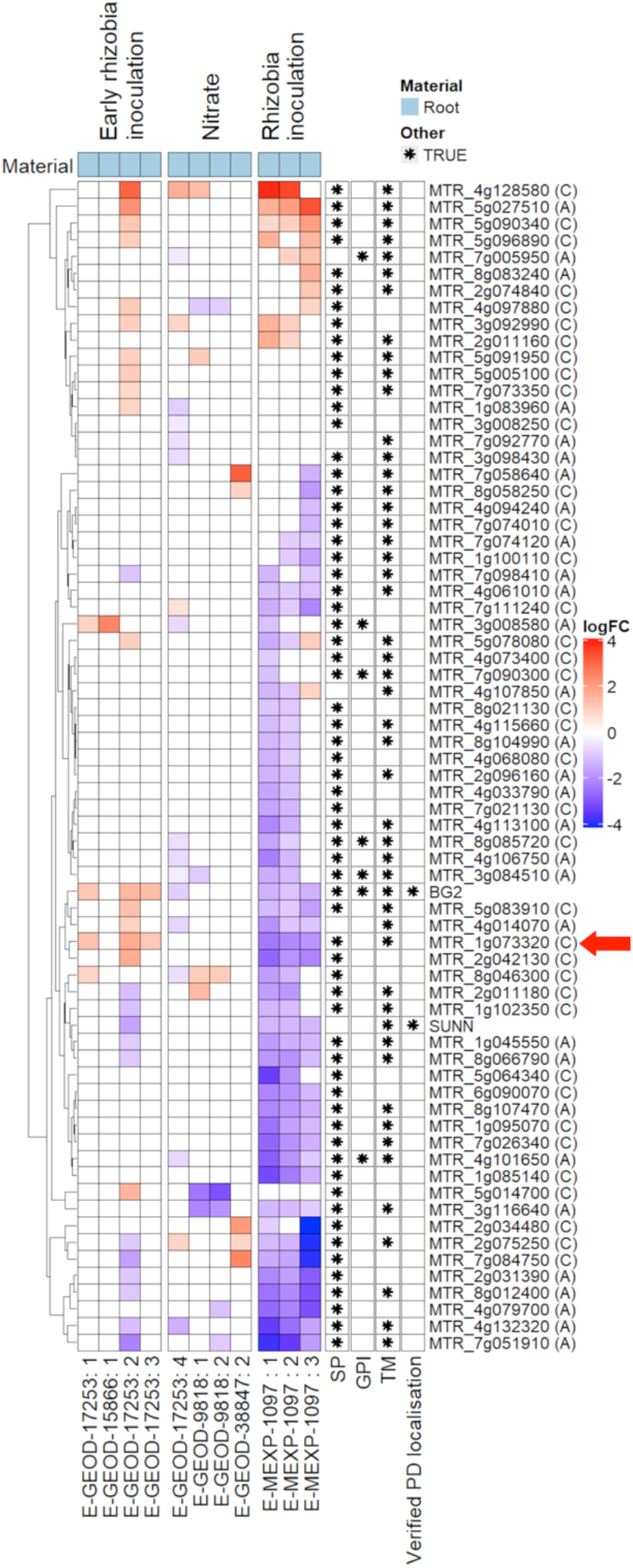
Expression analysis of *Medicago truncatula* PD candidates in rhizobia and nitrate transcriptomes. Gene expression analysis of PD candidates (lists A and C only) and verified PD genes BG2 and SUNN (see table S6). Differential gene expression (log_2_FC) was determined using public microarray data of experiments relating to nitrate and rhizobia inoculation in root tissue. Columns labels bottom= ArrayExpress accession codes followed by a reference number. See Supplemental table 8 for further information. (left dendrogram) Rows are ordered by hierarchical clustering. Asterisks in cells denote that gene has a membrane targeting feature or are verified. SP = Signal peptide, GPI = glycophosphatidylinositol anchor, TM = transmembrane domain. Red arrow indicates the position of MTR_1g073320, which is used in further analysis.

To established Medtr1g073320 localisation, C-terminal YFP fusions were created and introduced in *M. truncatula* roots using *A. rhizogenes*-mediated transformation. One week old transgenic roots were identified and counterstained with aniline blue which fluoresce in the presence of callose- a cell wall glucan component associated with PD. Confocal microscope images show Medtr1g073320-YFP in a punctate pattern on the cell periphery, co-localising with callose deposits indicating PD targeting (Figure 5).

**Figure 5:**
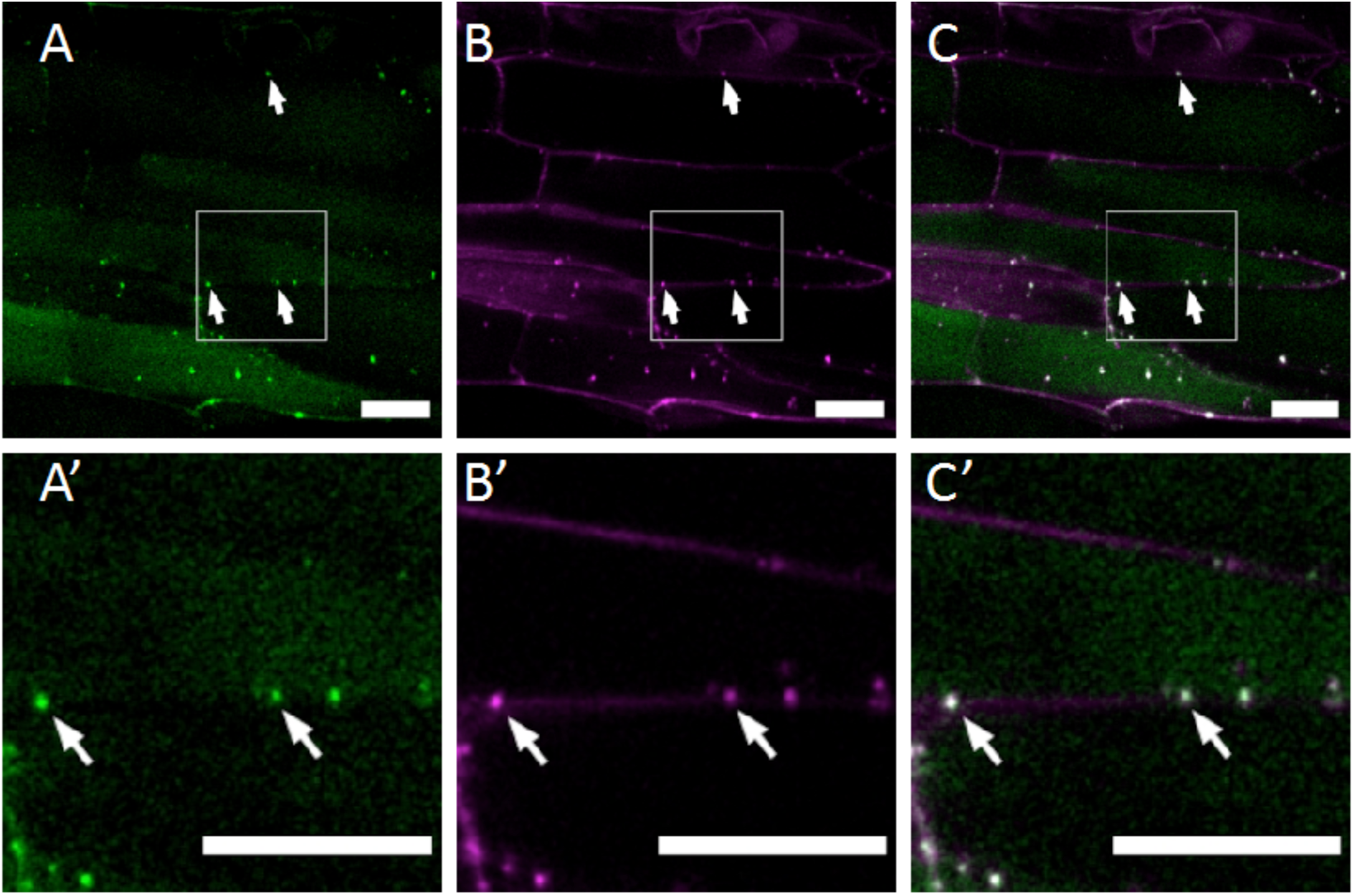
The *Medicago truncatula* protein Medtr1g073320 co-localizes with callose at PD microdomains. *Agrobacterium rhizogenes* transformed roots expressing Medtr1g073320 fused with YFP were counterstained with aniline blue to reveal callose deposition. Roots were imaged with a confocal microscope and (A) excitation laser 561 nm (green: Medtr1g073320-YFP) and (B) 405 nm (shown in magenta: aniline blue). Merged images are shown in C and co-localisation events highlighted with arrows. Bottom panels (A’-C’) are magnification of the area highlighted by the white box in the top panels. Scale bar = 20 μm.

Altogether, the results demonstrate how the pipeline, implemented in our custom build R script, can be used to predict the PD proteome in plant species where this information has not been experimentally derived, opening doors to the identification of novel PD proteins participating in the regulation of cell-to-cell communication.

## Discussion

Many aspects of PD function and regulation are not well understood despite the increasing evidence of the important role these structures play in plant signalling, organ development and response to physiological and environmental cues (Petit *et al.*, 2020; Vu *et al.*, 2020; Li *et al.*, 2021). Proteomic data is lacking for most plant species due to difficulties in isolating clean PD fractions and because association to PD domains can be quite dynamic, varying with environmental conditions and tissue types (Grison *et al.*, 2019; Hunter *et al.*, 2019; Cheval *et al.*, 2020). These factors limit our current knowledge of what proteins associate with PD and how their properties and function are controlled.

We created a pipeline that exploits the overlap in subfamily composition of existing PD proteomes to generate candidate PD genes (Figure 1). This allows the generation of *in silico* proteomes for 22 plant species which are represented in both PANTHER16 and in Ensembl Plant databases. The gene lists were classified according to their presence in one or more PD proteomes and according to predicted membrane targeting features overrepresented in verified PD proteins. Based on our categorisation, list A (subfamilies present in multiple proteomes and displaying predicted membrane targeting features) is more likely to contain PD localised proteins but genes in list C (subfamilies present in a single proteome) can not be excluded as only four PD proteomes are sequenced so far. Supporting our predictions, list A and C contained 36 out of the 60 proteins reported to target PD in *A. thaliana,* 46 if using families for *in silico* predictions (Table 1). The pipeline demonstrated to be effective in predicting the poplar proteome showing significant overlap with the experimentally determined PD enriched fraction (Fig. S3).

Combining proteomic with transcriptomic analysis we identified drought, osmotic and salinity stresses as conditions affecting the expression of clusters of genes identified in the Arabidopsis *in silico* proteomes. The diffusion in root meristems of GFP, expressed in companion cells under the SUC2 promoter, was restricted after continuous (4 days) and short (24h) exposure to 3% PEG and 75mM NaCl (Figure 3), suggesting that the expression of PD candidates is linked to the regulation of symplastic transport. We determined for the first time the *in silico* proteome of *M. truncatula* using PIP1 and exploited published microarrays to study gene expression during rhizobia infection and nodulation as conditions reported to affect symplasmic transport (Gaudioso-Pedraza *et al.*, 2018). This strategy led us to identify the receptor protein Medtr1g073320, which we confirmed localises at PD when expressed as fluorescent fusion in transgenic roots (Figure 5). Together our results validate the use of the PIP1 pipeline in combination with transcriptomic analysis for the identification of new PD proteins and conditions influencing symplasmic communication.

Our approach relies on subfamily annotation within the PANTHER database. For the 40 reference plant genomes included in PANTHER16, family/subfamily annotation coverage is between 60-95% depending on species (Mi *et al.*, 2021). For *A. thaliana* and poplar, the coverage is at 89% and 81% respectively but for *M. truncatula*, coverage is only 67%. Genes without a PANTHER classification will not be included in the output of the pipeline. We expect our approach may have less predictive power in monocots or non-angiosperms, as experimental proteomes are all from dicot species. It is unknown to what extend the composition of PD differs between monocots and dicots, especially as they differ in cell walls composition (Calderan-Rodrigues *et al.*, 2019). The experimental determination of more PD proteomes including representative monocot species and the identification of PD proteins by independent approaches will improve *in silico* predictions as these can be easily added as inputs to PIP1. Adding, as input data, subfamilies of PD proteins identified by different means can get around the limitations associated with the cell culture origin of experimental proteomes. Researchers already use sequence - domain analysis, phylogeny and transcriptomics to study families of PD proteins and to identify members expressed outside cell cultures (Thomas *et al.*, 2008; Simpson *et al.*, 2009; Gaudioso-Pedraza *et al.*, 2018). Our comparative meta-analysis provides a platform to systematically apply this approach enabling *in silico* predictions of whole PD proteome based in multiple instead of single experiments.

Applying the pipeline to *A. thaliana* or poplar generates a list of candidates larger than experimentally determined because expression in cell cultures is not a pre-requisite for gene identification. For example, the Arabidopsis *in silico* proteome contains BAM2, the Germin-like protein PDGLP1 and CALNEXIN2 (CNX2), which are not present in any of the experimental proteomes but localised at PD in independent studies (Fernandez-Calvino *et al.*, 2011; Liu *et al.*, 2017; Rosas-Diaz *et al.*, 2018; Brault *et al.*, 2019). The candidate lists also includes subfamilies/proteins lacking predicted membrane targeting features (SP, TM and GPI) (list B and D). Overlap between list B (subfamilies present in multiple proteomes) and the mobile proteome, recently reported in dodder parasitising *A. thaliana* (Liu *et al.*, 2020), suggest that these might be subfamilies of proteins identified while in transit via PD. List B might also include proteins with unusual membrane targeting mechanisms or with features poorly predicted by the available platforms. This is the case for REMORIN2, for example, which localises to PD independently of the secretory pathway and lacks a predicted SP, GPI or TM (Reymond *et al.*, 1996; Gronnier *et al.*, 2017; Huang *et al.*, 2019). Based on structural predictions and proteomic count, the lists serve as a tool to prioritise the screening, instead of rejecting, candidate PD proteins.

Besides the obvious reasons of obtaining *in silico* proteomes as a tool for the identification of new PD genes, we propose that transcriptomic analysis of the candidate lists can predict conditions affecting PD transport. Here, we showed that osmotic and salinity stresses modify the expression of clusters of PD candidates in Arabidopsis and that symplasmic transport of GFP is affected in these conditions. This aligns well with recent publications reporting changes in symplasmic transport and in PD protein localisation in response to salt and mannitol (Grison *et al.*, 2019; Hunter *et al.*, 2019). It is not clear if PD function is affected by changes in the localisation of PD proteins or by changes in turgor pressure or, more likely, to a combination of these (Hernández-Hernández *et al.*, 2019). The identification and phenotypic characterisation of mutants in PD genes and of relevant symplasmic mobile molecules are necessary to mechanistically understand the biological role of PD in the plant response to these abiotic stresses.

Our past research indicates that PD are regulated in response to rhizobia infection, a mechanism mediated by the degradation of the cell wall polysaccharide callose by the MtBG2 protein (Gaudioso-Pedraza *et al.*, 2018). PDLP regulation has been tightly linked to callose deposition in Arabidopsis (Lee *et al.*, 2011; Wang *et al.*, 2013; Caillaud *et al.*, 2014). Here, we identified Medtr1g073320 as a PD-located member of a subfamily of PDLP proteins (Thomas *et al.*, 2008) in *M. truncatula*. Expression tables suggest that Medtr1g073320 is co-regulated with SUNN and MtBG2 during rhizobia infection and nodulation. This finding opens doors for research on the mechanisms involving callose and PD regulation in the establishment of nitrogen fixation in legumes, an important process for sustainable agriculture.

To summarise, an R-based tool has been made available to integrate the data obtained from PD proteomes and enable the identification of new PD genes in a variety of plant species. Together with transcriptome analysis, the pipeline becomes a useful tool to identify conditions affecting PD composition and function. Our approach works with all plant species and genes with subfamilies annotated in both PANTHER16 and Ensembl Plant databases and will only expand as more new and updated UniProt Reference Proteomes are added (Drost & Paszkowski, 2017; Howe *et al.*, 2020; Mi *et al.*, 2021). The pipeline can help in prioritising the targets for validation from experimental proteomes and can predict PD proteins for species where PD proteomic information is not available. The pipeline is publicly accessible and can be easily modified by the user to add new sequenced proteomes and/or experimentally verified genes thus will serve as a long-lasting resource for the plant community.

## Acknowledgements

The authors would like to thank Hendrik Swiegers for his help in testing the pipeline and Dr. Eva Deinum and Dr. Michael Wilson for critical reading and help shaping the manuscript. This work was supported by UKRI Future Leader Fellowship (MR/T04263X/1), Leverhulme Trust Grant RPG-2016-136, a University of Leeds Gosden Studentship and a BBSRC DTP (BB/M011151/1).

## Author Contributions

P.K. designed and generated the pipeline, cross-referenced with published proteomics and databases and performed experiments and analysed data for *A. thaliana*. S.A. designed the R script for transcriptomic analysis; L.G. helped acquiring images and analysing data for *M. truncatula*. R.G.P designed and tested the construct for Medtr1g073320, generated transgenic roots and imaged them. Y.B.A. was in charge of conceptualisation, supervision, data curation, and writing- reviewing and editing the manuscript with the help of P.K. All authors discussed the results, edited and commented on the manuscript.

## Data Availability Statement

PIP1, the R-based pipeline described here, will be available for download along with associated database from GitHub (https://github.com/PhilPlantMan/PIP1) upon publication. Medtr1g073320 construct will be provided upon request.

## Notes

https://github.com/PhilPlantMan/PIP1

### Competing Interest Statement

The authors have declared no competing interest.

### Summary of Updates

Some writing mistakes have been corrected and 1 figure substituted for a table.

