## supplemental figures for "Comparative meta-proteomic analysis for the identification of novel plasmodesmata proteins and regulatory cues"

| |
|---|
| Supplemental table 1: PD proteins verified in *Arabidopsis thaliana* |
| A comprehensive list of PD-localising proteins in *A. thaliana* extracted from the literature at the time of publication. Gene ID, name, PANTHER subfamily and, when relevant, presence in the *in silico* or experimental proteome are provided. Citations reporting PD localisation are also included. |
| Supplemental table 2: Plant species listed in both Ensembl Plant databases and in PANTHER16 |
| The list contains all species compatible with the pipeline at the time of publication. Non compatible species may be used as a proteome input if genes are converted to orthologues of a compatible species. Compatible species are those listed in both PANTHER16 and Ensembl Plant databases. |
| Supplemental table 3: Subfamily counts in each PD proteome |
| The number of members of each subfamily were counted for each PD proteome. Every subfamily annotated in PANTHER16 that is represented in a PD proteome is listed here. |
| Supplemental table 4: *Arabidopsis thaliana in silico* proteome generated using PIP 1 pipeline |
| Candidate lists A-D are concatenated. Gene ID, candidate list where they appear, predicted structural features (SP, TM or GPI), PANTHER16 families and subfamilies names are shown. |
| Supplemental table 5: *Arabidopsis thaliana* genes in list A and C from families in which at least one member has verified PD localization |
| Candidate lists A and C are concatenated for genes of families with at least one member has PD verified localization. Gene ID, predicted structural features (SP, TM or GPI), PANTHER16 families and subfamilies names, record if gene, subfamily or family have a member localized at PD, proteome count where subfamily was identified and co-expression data are shown. |
| Supplemental table 6: *Medicago truncatula in silico* PD proteome generated using PIP1 |
| Candidate lists A-D are concatenated. Gene ID, candidate list where they appear, predicted structural features (SP, TM or GPI), PANTHER16 families and subfamilies names are shown. |
| Supplemental table 7: *Medicago truncatula* genes in list A and C from families in which at least one member has verified PD localization |
| Candidate lists A and C are concatenated for genes of families with at least one member has PD verified localization. Gene ID, predicted structural features (SP, TM or GPI), PANTHER16 families and subfamilies names, record if gene, subfamily or family have a member localized at PD and proteome count where subfamily was identified are shown. |
| Supplemental table 8: Metadata from published microarrays used in this study |
| The table contains information on the microarrays used for Figure 3, 5 and supplemental Fig. 4. Columns shows accession: experiment number, plant species, experiment title, material used, type/name, condition for differential expression and the corresponding links for more information of how each experiment was conducted. |

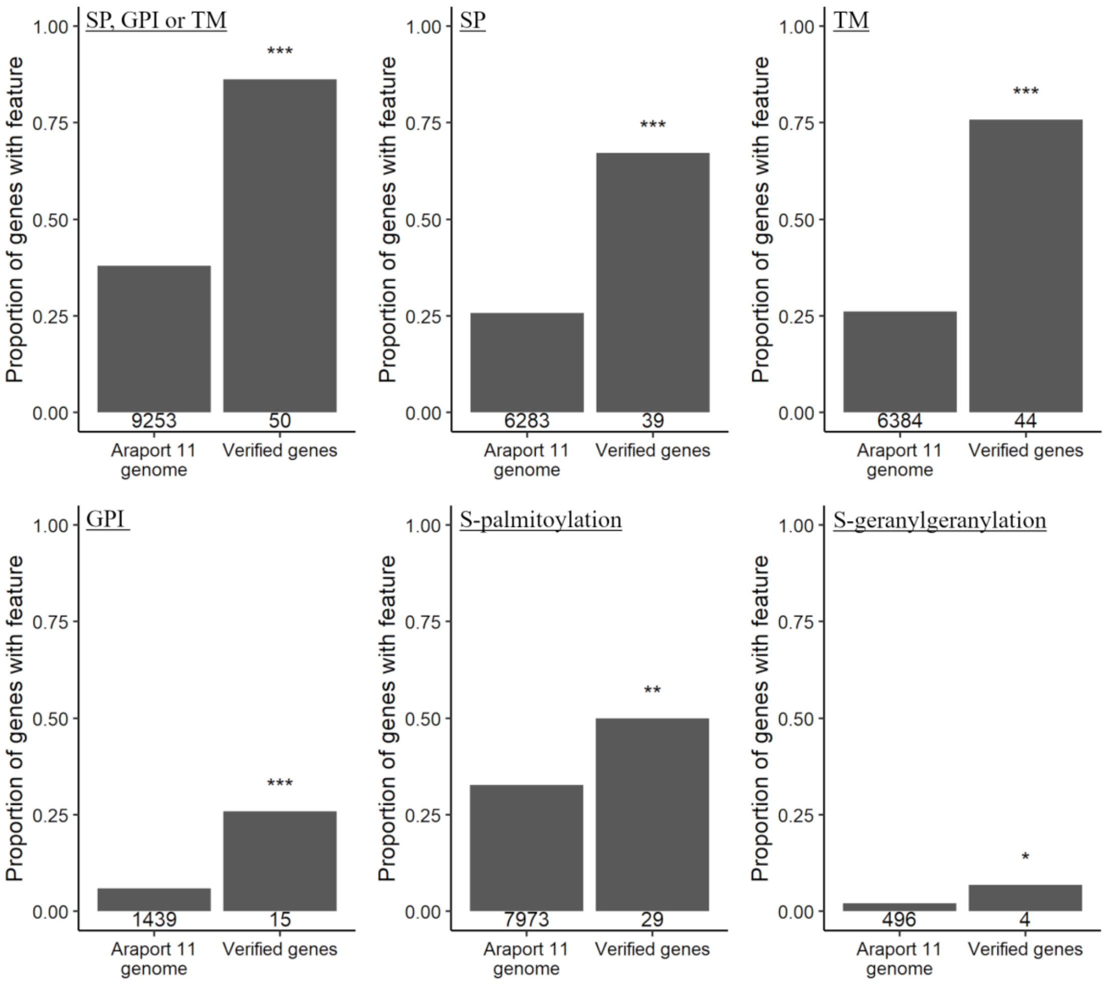

**Supplemental figure 1: Membrane targeting features encoded by verified PD genes.** Publicly available protein prediction tools were applied to peptide sequences encoded by the whole *Arabidopsis thaliana* genome. The proportion of verified PD genes with SP = Signal peptide, GPI = glycophosphatidylinositol anchor, TM = transmembrane domain, S-palmitoylation or S-geranylgeranylation was compared against the wider genome (Fisher's exact test, *: $p \leq 0.05$, **: $p \leq 0.01$, ***: $p \leq 0.001$). Number of proteins that are predicted to have the given feature are displayed underneath each column. Other protein features listed in the materials and methods were not found to be significantly overrepresented in verified PD genes (not shown). The specific s-palmitoylation prediction is 'cluster C' predicted using GPS-Palm (Ning et al., 2020). The specific S-geranylgeranylation prediction is 'cccxc' predicted using GPS-Palm (Xie et al., 2016). See materials and methods for further details and links used for predictions.

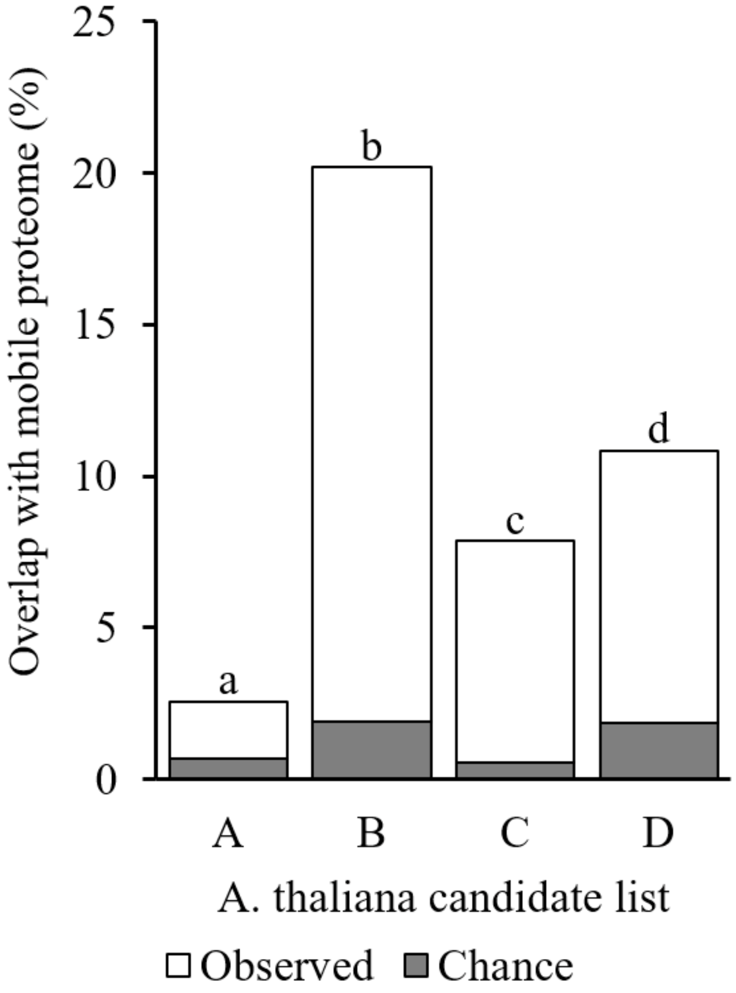

**Supplemental figure 2: *Arabidopsis thaliana* PD candidate lists are overrepresented in mobile proteins.** *A. thaliana* candidate proteome lists were compared to mobile proteins identified in *Cuscuta* parasitising *A. thaliana* (Liu et al., 2020) (mobile proteome). The size of the overlap for gene lists with the mobile proteome by chance was determined by bootstrap sampling of the whole genome (10,000 cycles, median % overlap given). Pairwise comparisons between observed overlaps of candidate lists and PD proteomes were made with Fisher's exact test ($p>0.05$, holm adjusted, bars with differing letters are significantly different).

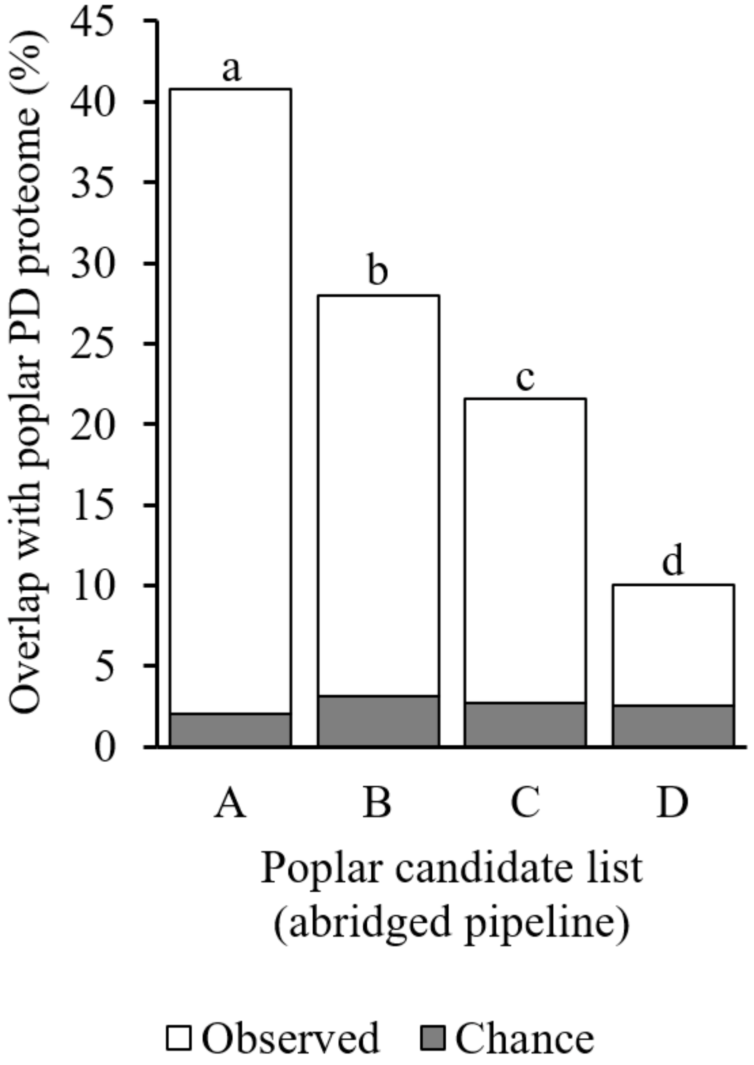

**Supplemental figure 3: Overlap between *in silico* and experimentally determined proteome is dependent on candidate list classification.** Poplar (*Populus trichocarpa***)** candidate PD genes lists A-D were generated using an abridged PD candidate pipeline that omitted the poplar PD proteome (Leijon et al., 2018) as an input. The size of the overlap for gene lists with the PD proteome by chance was determined by bootstrap sampling of the whole poplar genome (10,000 cycles, median % overlap given). Pairwise comparisons between observed overlaps of candidate lists and PD proteomes were made with Fisher's exact test ($p>0.05$, holm adjusted, bars with differing letters are significantly different).

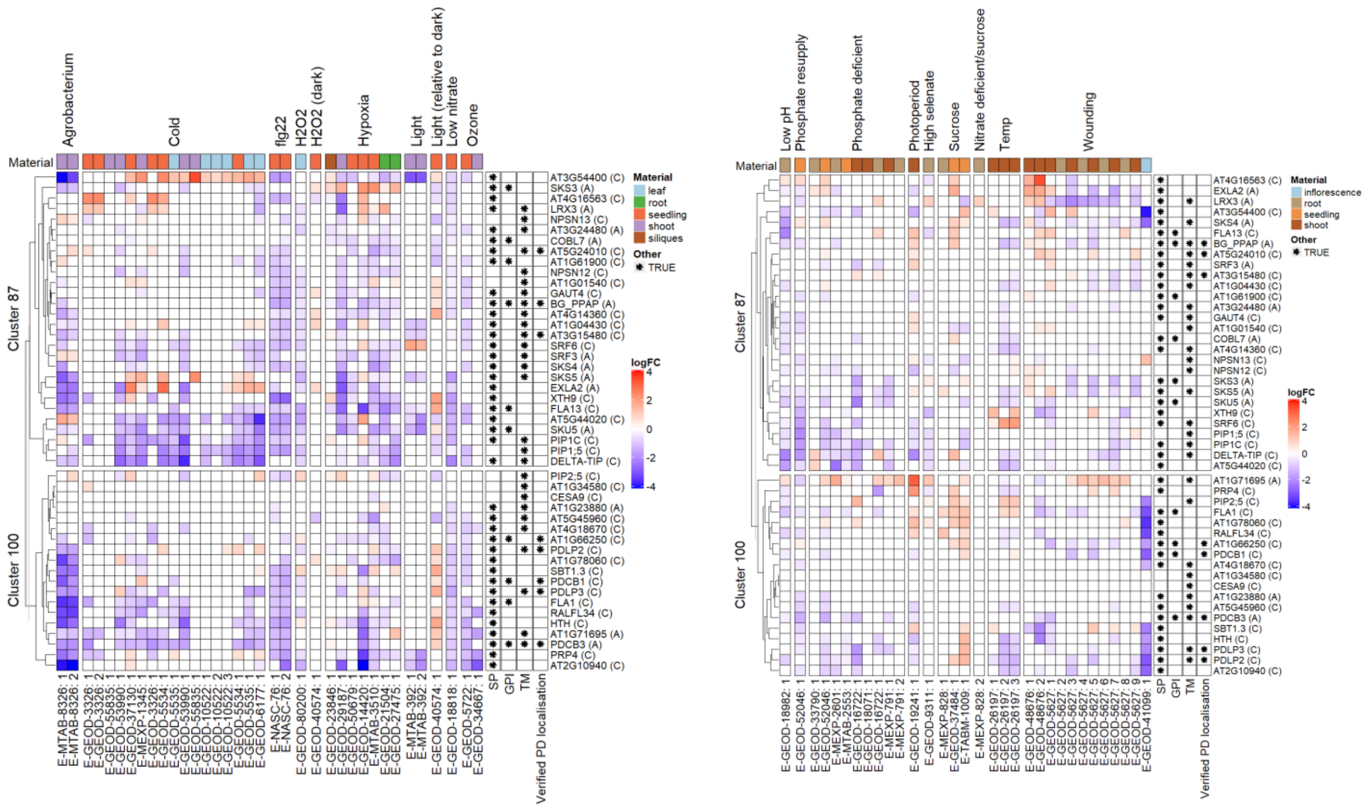

**Supplemental figure 4: Expression analysis of *Arabidopsis thaliana* PD candidates and PD verified genes in abiotic and biotic stress transcriptomes.** Gene expression analysis of verified subfamily members and candidate genes from clusters 87 and 100. Differential Gene expression ($log_2FC$) was determined from public microarray data. This figure complements data from the experiments presented in Figure 3. Columns labels = ArrayExpress accession codes followed by a reference number. See Supplemental table 6 for further description of the experiments. (left) Rows are grouped by cluster and then (dendrogram) ordered by hierarchical clustering. Asterisks in cells denote predicted features: SP = Signal peptide, GPI = glycosylphosphatidylinositol anchor, TM = transmembrane domain. Cell colour above each column = plant material sampled.
